# Cultural diffusion dynamics depend on behavioural production rules

**DOI:** 10.1101/2021.12.22.473828

**Authors:** Michael Chimento, Brendan J. Barrett, Anne Kandler, Lucy M. Aplin

**Affiliations:** Cognitive and Cultural Ecology Research Group, Max Planck Institute of Animal Behavior; Am Obstberg 1, 78315 Radolfzell, Germany; Centre for the Advanced Study of Collective Behaviour, University of Konstanz; Universitätsstraße 10, 78464 Konstanz, Germany; Department of Biology, University of Konstanz; Universitätsstraße 10, 78464 Konstanz, Germany; Department for the Ecology of Animal Societies, Max Planck Institute of Animal Behavior; Am Obstberg 1, 78315 Radolfzell, Germany; Department of Human Behavior, Ecology and Culture, Max Planck Institute for Evolutionary Anthropology; Deutscher Platz 6 04103 Leipzig, Germany; Division of Ecology and Evolution, Research School of Biology, The Australian National University, 46 Sullivan Creek Road, Canberra, ACT 2600, Australia

**Keywords:** cultural evolution, social learning, agent based model, network-based diffusion analysis, experience weighted attraction models

## Abstract

Culture is an outcome of both the acquisition of knowledge about behaviour through social transmission, and its subsequent production by individuals. Acquisition and production are often discussed interchangeably or modeled separately, yet to date, no study has accounted for both processes and explored their interaction. We present a generative model that integrates the two to explore how variation in production rules might shape cultural diffusion dynamics. Agents make behavioural choices that change as they learn from their productions. Their repertoires also change over time, and the social transmission of behaviours depends on their frequency. We diffuse a novel behaviour through social networks across a large parameter space to demonstrate how individual-level behavioural production rules influence population-level diffusion dynamics. We then investigate how linking transmission and production might affect the performance of two commonly used inferential models for social learning; Network-based Diffusion Analysis, and Experienced Weighted Attraction models. Clarifying the distinction between acquisition and production yields predictions for how production influences diffusion that are generalisable across species, and has consequences for how inferential methods are applied to empirical data. Our model illuminates the differences between social learning and social influence, demonstrates the overlooked role of reinforcement learning in cultural diffusions, and allows for clearer discussions about social learning strategies.

## 1 Introduction

Cultural evolution, or the changes over time in distributions of the types, forms or functions of socially-learned traits, provides a secondary inheritance system that potentially enables rapid adaptive plasticity [1, 2, 3, 4]. Social learning, the process that underlies cultural evolution, has a well accepted definition of learning that occurs through observations of the behaviour, or interactions with the products of behaviour, of others [5]. It is perhaps well accepted due to its ambiguity, as it encompasses many different phenomena. In particular, social learning may refer to 1) the learning of a novel skill from conspecifics, such as a cockatoo learning to open bins from associates [6], or 2) the influence that social information exerts upon behavioural choice, such as when a stickleback fish chooses feeders surrounded by more conspecifics [7]. In the first case, learning describes an event of acquisition of knowledge about a new behaviour as a result of social transmission [8]. In the second case, learning describes the changes in weights given to known behaviours when producing a behaviour, due to social influence. This ambiguity is further complicated by the recursive relationship between these two phenomena. Acquisition requires observable productions, captured in the definition of social transmission: when a behaviour is produced by an individual, it exerts a positive causal influence on the rate at which that individual’s associates acquire the same behaviour [8]. The conceptual distinction of origin (acquisition) and maintenance (production) of behaviour was previously highlighted by Galef [9] in the debate over why social learning is adaptive. Galef identified one critical cognitive factor that influences production directly: reinforcement learning, which he hypothesized influenced the maintenance or abandonment of cultural traits [9]. However, little is understood about how cultural dynamics are influenced by variation in such learning rules.

The breadth of the term “social learning”, and the circular relationship between acquisition and production, can also lead to ambiguity over the precise target of causal factors that influence cultural evolution. Generally, acquisition depends on cognitive [10, 11, 12], social [8, 13, 14], and environmental factors [15, 16]. These factors can also influence realized production behaviour [16, 17], *but not necessarily in the same way as acquisition*. There may also be imprecision over the target of social learning biases or strategies—do they act upon the process of acquisition or production, or both? This has lead to potential for miscommunication, for example in the long-standing debates over definitions of conformity [18]. Here, we aim to clarify the relationship between acquisition and production. We develop a simulation framework that treats these concepts separately, but permits their interaction via a process of social transmission [8]. Our mathematical model builds upon Galef’s verbal model and provides new predictions for how changes to learning rules of production influence diffusion dynamics, and offers clarity in discussions of the targets of social learning biases, strategies or constraints.

Acquisition and production have been modelled separately as approaches to two central goals of empirical studies of animal culture. One goal has been to identify when social transmission is responsible for the diffusion, or spread, of a behaviour through a population, as opposed to asocial innovation. A primary determinant of social transmission is the opportunity for naïve individuals to learn from others, represented by their social association. This correlation between association and social transmission is at the core of network-based diffusion analysis (NBDA) [8, 14, 19], which has been used to identify social learning (*sensu* social transmission) in birds [6, 20, 21, 22], fish [23, 24], cetaceans [25, 26, 27], and primates [28, 29, 30, 31]. Another goal is the identification of social influence, or how individuals integrate social information during decision-making between behaviours. This has been studied using experience weighted attraction (EWA) models—an extended reinforcement learning model originally developed to understand how behavioural choices change over time in economic games [10, 32, 33]. Due to their flexibility, EWA models are a popular method to identify social learning (*sensu* social influence on production). EWA has been used to study social learning strategies, both theoretically [34, 35] in humans [10, 36, 37, 38] and non-human animals [4, 39, 40, 41].

As with any model-based inferential framework, both NBDA and EWA have short-comings. NBDA assumes that the order of acquisition is correlated to network structure in cases of social transmission. However, acquisition of knowledge is not directly observable without behavioural output. Thus, NBDA assumes that the order of first observed production of a behaviour *is equivalent* to the order of acquisition. Divergences between acquisition of knowledge and first-productions might spell trouble for this assumption. Secondly, NBDA analyses do not usually account for behavioural frequency information (although recommended [14, 42], see [20, 43, 44]). Regarding EWA, its formalization requires the definition of a set repertoire of behaviours which an individual may produce. To date, implementations have not accounted for differences in repertoire size over time or between individuals, and thus are restricted to situations where there are no innovations or diffusion, as individuals must have homogeneous repertoires. In summary, a model of acquisition is not complete without a model of production, and vice versa. To describe dynamics of cultural change with some generality, we must simultaneously account for potential acquisition of novel behaviour, and the complex decision-making that contributes to the maintenance of behaviour. Yet, beyond specialized models of language evolution [45, 46, 47], no general, species-agnostic model of cultural evolution exists in which both acquisition and production are connected.

Here, we develop a simulation model that unites both processes by adapting the dynamics of NBDA and EWA frameworks. First, we explore how changes to production rules might affect the diffusion of a novel behaviour through networks of agents. We uncover new relationships between production and diffusion dynamics, and find that certain production rules result in more or less divergent orders between acquisition and first-production events. However, inferential NBDA models are blind to such divergences, and EWA models are blind to any heterogeneity in repertoire. To understand the consequences of this for the practical application of inferential methods, we then apply inferential models of NBDA and EWA to data generated by our model. We generate agents’ acquisition and production times, and behavioural output under known underlying mechanisms, e.g. social transmission or asocial innovation, and feed these data into inferential NBDA and EWA models. We find that linking production and acquisition limits the inferential power of these two popular methods. Finally, we discuss how our framework helps clarify broader discussions of social learning biases and strategies.

## 2 Methods

Our model simulated both the diffusion of a novel behaviour through a population, and the agents’ decisions of which behaviours to produce throughout the diffusion. Throughout the paper we used “diffusion” to refer to the spread of knowledge of the novel behaviour through a population. “Acquisition” was when an agent learned that they may produce it. “First-production” was when an agent first performed it. Acquisition and production were implemented as two sub-models that were mathematically consistent with prior applications of inferential NBDA [14] and EWA [10] frameworks, respectively. Some conventional notation has been altered be more accessible. Inferential NBDA statistically tests for acquisition of behaviour via social transmission against a null hypothesis of asocial innovation. It estimates the plausibility of social transmission by accounting for association with knowledgeable individuals, and orders or times of acquisition by naïve individuals. Inferential EWA is a dynamic learning model that uses time-series behavioural data to estimate how individuals balance social and personal information, as well as recent and past information, in decision-making. While there are many ways to formulate either model of acquisition or production, we have chosen to follow these specific frameworks to robustly evaluate how linking acquisition and production might affect the inferential value of either model.

Importantly, we transformed these inferential models into a generative model. We used their assumed dynamics to produce probabilities of acquisition or production of a novel behaviour—generating data using known parameters, rather than inferring parameters from observed data. We connected the two sub-models, allowing them to influence one another. The sub-model of acquisition determined agents’ repertoires, i.e. behaviours able to be produced, while the sub-model of production determined which behaviour was chosen for production from their repertoires in each time step, providing behavioural frequency information. Those frequencies then influenced the acquisition probability of a novel behaviour by naive agents in the following time-step. Thus, our model’s output was the cultural dynamic generated by linked processes of acquisition and production, conditioned on specific parameter values of either sub-model. We also explored the dynamics of the sub-models of acquisition and production in isolation (detailed in supplementary **Text S1**).

We assumed a population of constant size *N* = 24. Agents were situated within a social association network that constrained observation and acquisition opportunities to connected nodes. We used a random regular network architecture, with fixed degree (*k* = 6) associates for each agent (**Figure S1A**). Random regular architecture was chosen because degree was important to standardize between agents—it directly affected the dynamics of acquisition (detailed in **Equation** (2)). Edge weights were set to 1 to eliminate stochastic variation arising from differences in association. We explored other network architectures (**Text S2**), but the primary focus of this paper was to understand how learning rules governing behavioural production, rather than social structure, influenced diffusion dynamics.

At the beginning of a simulation, all agents initially had a repertoire of one behaviour, *a*, interpreted as an established tradition that had already diffused through the population. To remove stochasticity from differences in innovation timing, one randomly selected “seed” agent also had knowledge of a novel behaviour *b*. Both behaviours obtained an equal payoff (*π*_*a*_ = *π*_*b*_ = 1). Each time-step, agents had the opportunity to expand their repertoire by acquiring behaviour *b*, if it had been produced by at least one of their associates (see **Section 2.1**). Additionally, agents chose one behaviour from their repertoire to produce. Their choice was influenced by past personal experiences of payoffs, as well as social information from their neighbours (see **Section 2.2**). Here reinforcement learning played an important role: when an agent acquired knowledge of behaviour *b*, its expected payoff was initially 0. Expectations were learned only through individual experience of producing the behaviour (see Equation (4)). This cycle of potential acquisition and production repeated until the novel behaviour *b* had been acquired and produced at least once by each agent.

### 2.1 Sub-model of behavioural acquisition

Each time-step, agents had the opportunity to acquire knowledge of novel behaviour *b* through social transmission. The probability of acquisition depended on an agent’s associates and their behavioural productions, a rate of social transmission, and a rate of asocial innovation. We estimated the probability of acquisition of behaviour *b* for each naïve individual *i*, i.e. individuals that only possessed behaviour *a*, at each time-step *t* by calculating:

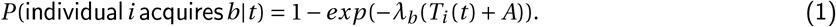

The baseline learning rate (*λ*_*b*_) represented how easily the novel behaviour was learned, either socially or asocially. *A* represented the presence of asocial innovation, and could take a value of 0 or 1.

The transmission function *T*_*i*_ (*t*) recorded the relative usage of behaviour *b* of all knowledgeable associates of agent *i* over a memory window of *m* time-steps. The transmission function *T*_*i*_ (*t*) was defined by

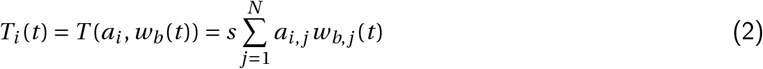

whereby the sum was taken over agent *i*’s associates as *a*_*i, j*_ = 1 if *i* and *j* were connected in the association network and *a*_*i, j*_ = 0 otherwise. *w*_*b, j*_ (*t*) defined the transmission weight of agent *j* representing the proportion of time this agent produced behaviour *b* within the last *m* time-steps

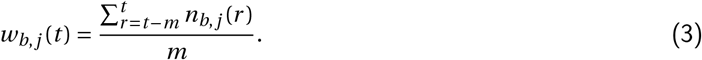

We set *n*_*b, j*_ (*r*) = 1 if agent *j* produced *b* at time step *r* ; otherwise *n*_*b, j*_ (*r*) = 0. Lastly, the sum was multiplied by *s*, the rate of social transmission. Higher values of *s* resulted in increased probability of transmission per knowledgeable associate, relative to the asocial innovation.

In summary, the transmission weight function was the nexus of our sub-models of acquisition and production, as illustrated by **Figure 1**. It allowed behavioural frequencies, derived from the production sub-model detailed in the following section, to influence transmission probabilities calculated from the acquisition sub-model.

**Figure 1:**
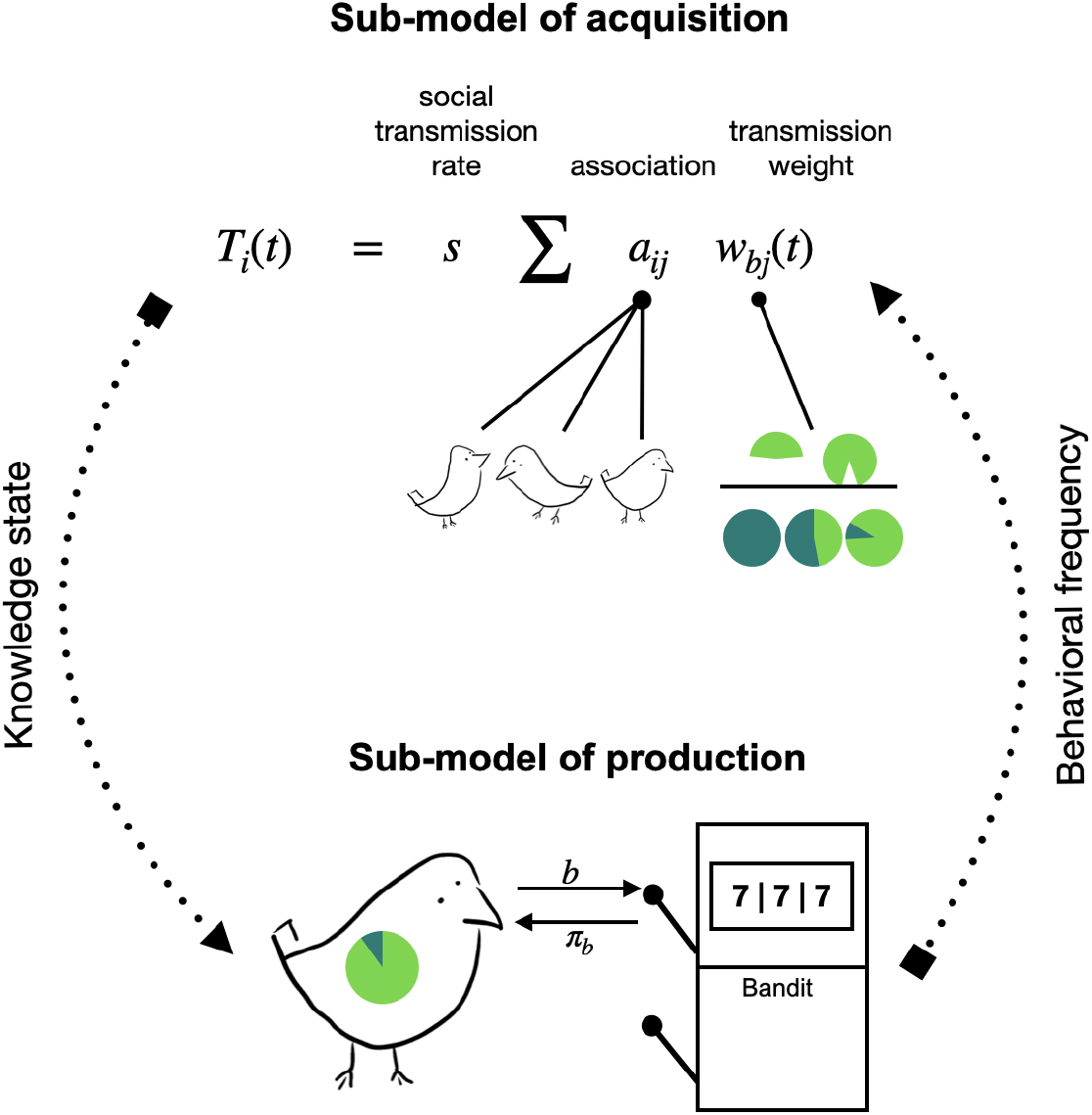
Model schematic. Each time-step, naive agents had the opportunity to acquire knowledge about novel behaviour *b* from their associates. Acquisition was conditioned on association and the frequencies that associates produced *b*, as well as a rate of social transmission, as defined by the transmission function *T*_*i*_. If an agent acquired *b*, they could then produce *b* using the sub-model of production. Production probabilities changed over time, depending on personal experience and social information. behavioural frequency information from the production sub-model then informed acquisition probabilities in the following time-step.

### 2.2 Sub-model of behavioural production

Each time-step, each agent *i* produced a behaviour *k* from its repertoire *Z*_*i*_ (consisting of either {*a*} only or {*a, b*}) with probability *P*_*i, k*_ (*t*) as follows. Agents updated their expected payoffs given their personal experience of producing a behaviour at *t* − 1 (**Equation** (4)), which influenced the probability of producing either behaviour given personal information only (**Equation** (5)). However, agents also considered social influence in their choice using a record of the observed productions of its associates (**Equation** (6)). Changes in personal and social information potentially led to a changed production probability at time *t* (**Equation** (7)). 4 parameters governed how these values changed with experience: recent experience bias (*ρ*), a risk-appetite bias (*α*), a social information bias (*σ*) and frequency-dependent production bias (*χ*).

First, agent *i* calculated the expected payoff (*E*_*i, k*_ (*t*)) for any behaviour *k* in its repertoire using

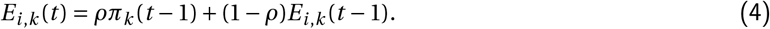

*π*_*k*_ (*t* − 1) described the payoff obtained from behaviour *k* in the last time-step: if behaviour *k* has been produced *π*_*k*_ (*t* − 1) = 1 otherwise *π*_*k*_ (*t* − 1) = 0. This assumption allowed for information loss over time. In the first time-step, agents were initialized with *E*_*i, a*_ (1) = 1. Once an agent acquired the novel behaviour, *E*_*i, b*_ (*t*) = 0. For behaviour *k, max*(*E*_*i, k*_) = *π*_*k*_, ∀ *i*, meaning expected payoffs could never exceed the true payoff. *ρ* determined the importance of the most recently received payoff *vis-a-vis* previously experienced payoffs.

In the second step, agent *i* converted its expected payoff from the behaviours in its repertoire *Z*_*i*_ into a probability using a softmax rule

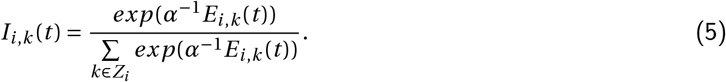

The parameter *α* controlled the sensitivity towards differences between expected payoffs: low values resulted in an almost deterministic choice of the highest expected payoff behaviour (risk-aversion), whereas high values lead to choices almost independent from expected payoffs. **Equation** (5) assigned probabilities to behaviours even when their expected payoff was 0, providing the possibility of production without prior experience.

Next, agent *i* evaluated its social information by counting how many times its associates had produced any of the behaviours in its repertoire within the memory window comprised of the last *m* time steps. This was modulated by the frequency dependent production bias parameter *χ*, which determined the strength of frequency dependent influence on the agent

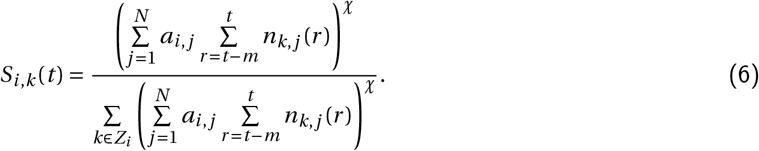

Agents only observed behaviours which were in their own repertoire. If an agent did not know how to produce a behaviour, its observation could not influence an agent’s choice in that time-step. However, this information did influence the potential acquisition of this behaviour as described in **Section 2.1**.

Lastly, agent *i* combined their personally experienced payoffs and socially observed behaviours to generate probabilities of producing the behaviours in its repertoire

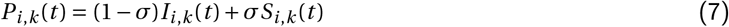

where *σ* described the preference for individual and social information. Values of *σ* close to 0 would almost neglect social information while values close to 1 would almost neglect individual information.

### 2.3 Conditions

In order to understand how production rules influenced diffusion dynamics, we explored diffusions across populations under a variety of parameter settings, summarized in **Table** 1. Each combination of these parameters formed one constellation. For each constellation, we ran 100 simulations.

In **Section 3.1** we used the following parameter constellation as reference

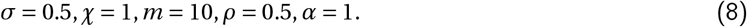

We focused on results from the combinations of constellations described by **Table 1**, using a sensitivity analysis to explore the effect of each parameter while holding all others at reference. Intermediary and extended values of these parameters were also evaluated (e.g. *σ* ∈ {.125,.375,.625,.875}), although they showed no unexpected patterns and were thus excluded from the paper for concision. The effect of *ρ* on the evolution of expected payoffs in **Equation** (4) is non-linear (i.e. the difference in effect between *ρ* = .013 and *ρ* = .014 is magnified compared to the difference in effect between *ρ* = .79 and *ρ* = .80)), thus we tested 3 different orders of magnitude, rather than differences in linear magnitude. We also varied parameters related to the acquisition sub-model in the supplementary material (see **Text S2** for network structure, *λ*_*b*_, *s* and *A* values) but have excluded these from the main text, as the effect of network structure and social transmission on diffusion is a well-explored subject [48, 49, 50]. The baseline learning rate was set to *λ*_*b*_ = 0.05, the social transmission rate was set to *s* = 5. In order to isolate the effect of variation in production rules on diffusion via social transmission, and eliminate any variation due to random innovation events, the asocial innovation parameter was set to *A* = 0 for **Section 3.1**. However, its unrealistic to expect that no independent innovation occurs in natural diffusions. Thus, *A* = 1 in simulations used to test the performance of inferential models (see **Section 2.5**).

**Table 1:**
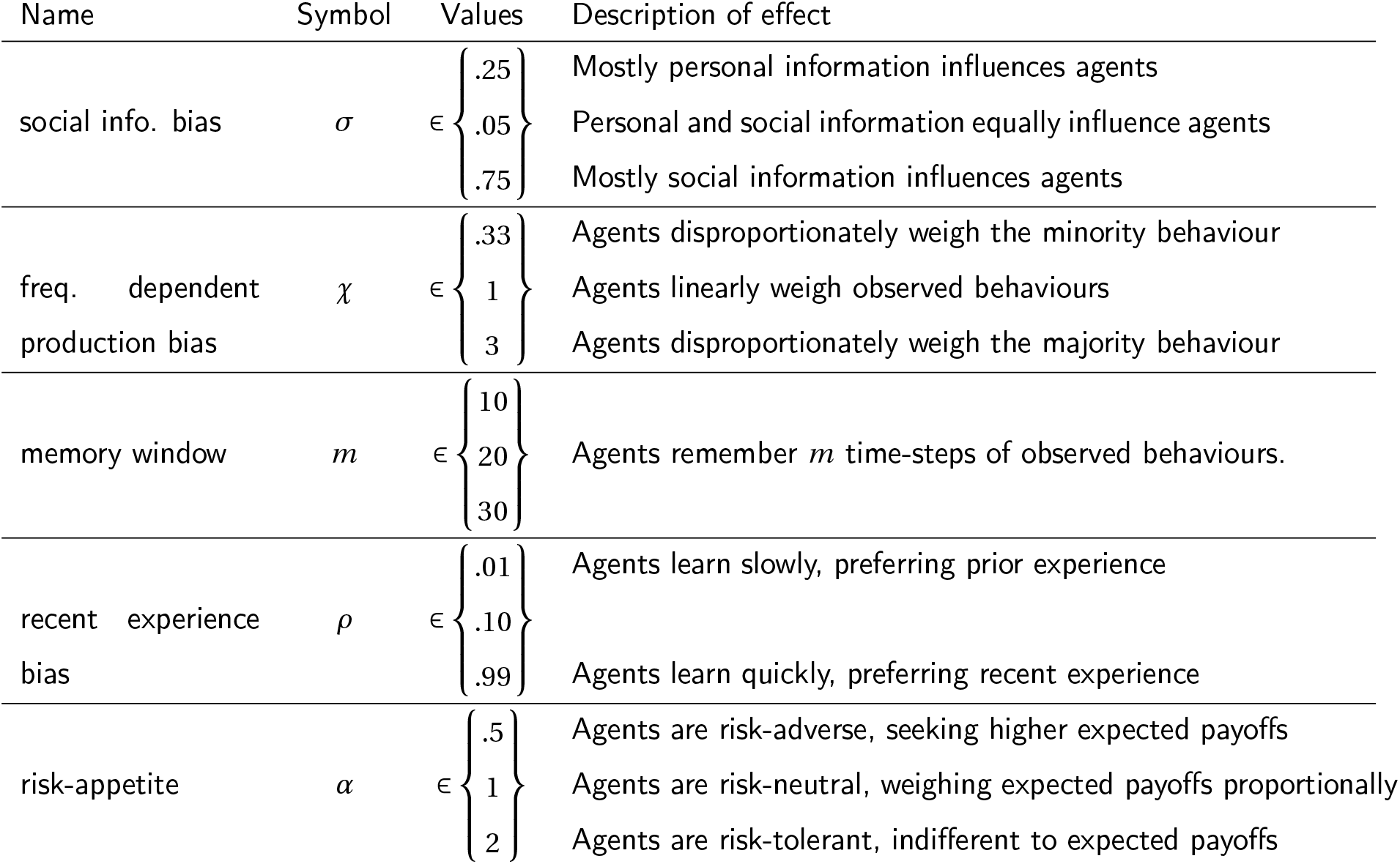
Summary of tested parameters for the production sub-model. Mathematical symbol, name, values tested, and short description of these values.

### 2.4 Measurements

To describe the dynamics of the diffusion of variant *b* through the population we calculate three quantities: tempo, divergence and delay. To quantify diffusion tempo, we recorded the time-steps at which each agent acquired the novel behaviour and first produced the novel behaviour. We also recorded behavioural frequencies every time-step. From these data we derived time-to-diffusion (TTD), i.e. the time until the whole population had acquired behaviour *b*, intervals between each acquisition event, time-to-first-production (TTFP), i.e. the time until the whole population has produced behaviour *b* at least once, vectors of order of acquisition (*o*_*a*_), order of first-production (*o*_*p*_), time of acquisition (*t*_*a*_) and time of first-production (*t*_*p*_). Vectors *o*_*a*_ and *o*_*p*_ contained the position of each agent in the acquisition sequence and first-production sequence, respectively. Vectors *t*_*a*_ and *t*_*p*_ contained the time of acquisition and time of first-production by each agent, respectively.

To quantify divergence in the orders of acquisition and first-production, we calculated the mean Manhattan distance

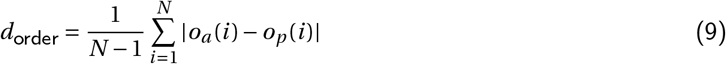

representing the mean difference between acquisition and first-production position in the population (excluding the first-production of the seed agent, i.e. the agent at position 1, thus *N* − 1, as *o*_*a*_ (1) − *o*_*p*_ (1) = 0 per definition). Additionally, we calculated the proportion of the population who obtained production positions that differed from their acquisition positions.

To quantify the delay between the acquisition and first-production timing, we used vectors *t*_*a*_ and *t*_*p*_ and calculated the mean difference between the times of acquisition and first-production

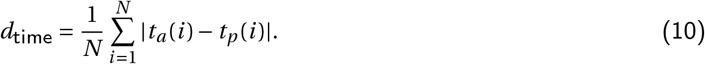

For each measurement we reported means and 92% percentile intervals in brackets.

### 2.5 Testing performance of NBDA and EWA on generated data

Lastly, we explored how NBDA and EWA might perform on data generated by our model. NBDA assumes that the order of first-production is equivalent to that of acquisition. Yet, when acquisition was conditioned on production (as in our model) these orders could diverge. To test how this might affect the inference of the underlying generative process (i.e. social transmission or asocial innovation), we first created four data-sets: two sets of “ideal” data generated using the pure NBDA dynamic with which we expected NBDA to provide strong support for the correct generative process, and two data-sets of more realistic data that violated NBDA’s assumption that orders of acquisition and first-production were identical. To create ideal data, we recorded time-of-acquisition data, generated by either asocial innovation only (*A* = 1, *s* = 0), or social transmission (*A* = 1, *s* = 5) where the transmission weight was *w*_*j, b*_ (*t*) = 1 if associate *j* was knowledgeable, otherwise 0, removing the influence of production on transmission. To create realistic data, we recorded time-of-first-production data, generated by either social transmission using transmission weights as we have defined them, or asocial innovation only. For each of these four scenarios, we simulated 10 diffusions at each parameter constellation (**Table 1**).

We then ran inferential TADAc NBDA models [14] on each simulation’s data. For simulations with social transmission, the seed agent was included as a demonstrator. For realistic data, we included a transmission weight in the inferential models, defined as a production rate of the novel behaviour for each agent (total productions of *b* divided by time-steps knowledgeable). Using recommended inferential steps, we ran both a social and asocial TADAc models to determine support for social transmission [8]. We compared AICc scores between the two models to determine relative support for social or asocial innovation [8]. Support for social transmission was defined as Δ*AICc* > 0, where Δ*AICc* = *AICc*_*asocial*_ − *AICc*_*social*_. We reported the median of all Δ*AICc*s per condition to get an idea of what the average support for the generative process might be.

We used a similar strategy to evaluate the performance of EWA analyses. We estimated only recent experience bias (*ρ*) and social information bias (*σ*) under three scenarios: 1) homogeneous repertoires and no diffusion, 2) heterogeneous repertoires with only social transmission (*A* = 0, *s* = 5), and 3) heterogeneous repertoires with only asocial innovation (*A* = 1, *s* = 0). Scenario 1 was a proof of concept with ideal data generated by using the pure EWA dynamic, and the EWA model was expected to recover accurate estimates. Scenarios 2 and 3 explored how EWA performed on data collected from populations with heterogeneous repertoires, when behaviours were actively diffusing or being innovated. For each scenario, we performed 10 simulations for each combination of *ρ* and *σ* values ({0.25, 0.5, 0.75} for both). All other parameters were set to reference.

For scenario 1, we recorded behavioural productions for 300 time-steps to provide sufficient power. In scenarios 2 and 3, each simulation ran for twice as long as its TTD (social transmission: range [65 − 199], asocial innovation: [89, 235] time-steps), thus recording equal numbers of observations both during diffusion, and after all agents acquired knowledge of the novel behaviour. We then fit inferential models to the production data from each simulation from these three scenarios using Hamiltonian Markov Chain Monte Carlo (MCMC). Models were run using 3 chains, 2000 iterations, with 1000 warm-up iterations, with all estimates based on over 1000 effective samples from the posterior. All models were fit using R v. 4.0.2 [51] with Stan v. 2.27 [52] via Rstan v. 2.21.2 [53]. Good model convergence was confirmed from evaluating rank histograms and ensuring parameters had Gelman-Rubin’s statistic 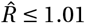 ≤ 1.01 [54, 55]. Additionally, we plotted priors against the posteriors to assess how well the models identified parameters. For each parameter, we report mean and 92% highest posterior density interval (HPDI), or the narrowest interval contains 92% of posterior samples.

## 3 Results

Simulations with linked acquisition and production sub-models generated the dynamics depicted in **Figure 2A**. At the reference parameter constellation (**Equation** (8), henceforth reference), knowledge of the novel behaviour diffused to all individuals in 54.22[37.96, 76.28] time-steps (mean TTD [92% PI]). The first-production curve lagged behind the knowledge acquisition curve, with all agents producing the novel behaviour at least once in 61.11 [44.96, 85.04] time-steps (mean TTFP). Agents retained a higher expected payoff for the established behaviour and still preferred its usage by the time full diffusion was reached. However, the behaviours trended towards an equilibrium production ratio of 1 : 1, as both yielded equivalent payoffs.

**Figure 2:**
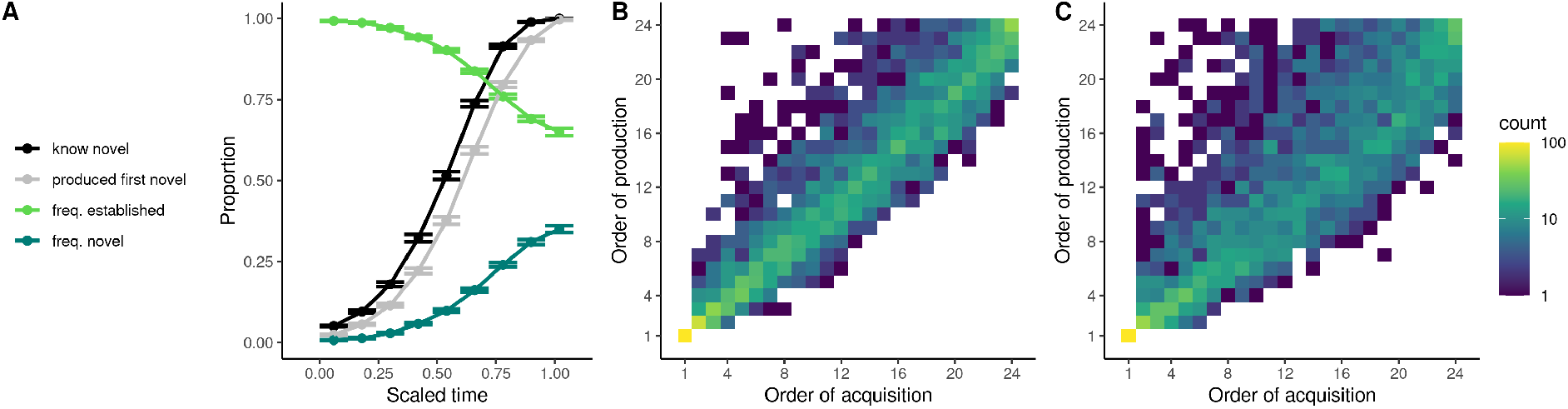
Acquisition and production dynamics throughout diffusion. Data presented from 100 simulations at reference setting. **A)** Average knowledge acquisition curve of novel behaviour (black), first-production curve of the novel behaviour (grey), and behavioural frequency data of the established behaviour (light green) and the novel behaviour (dark green). **B)** Density plot comparing the order in which the novel behaviour entered an agent’s repertoire (*x*-axis) and the order in which that agent first produced the novel behaviour (*y* -axis) at reference setting. Each count is one agent from one simulation. Measuring the order of first-production usually did not reflect the order of acquisition, and only about 25% of all agents produced the novel behaviour in the same order that they acquired it. **C)** Density plot for the parameter setting that obtained the most divergence (83.2% divergent agents; *σ* = 0.25, *χ* = 3, *ρ* = .99, *α* = .5, *m* = 10).

Importantly, production did not simply lag behind knowledge acquisition—we observed divergences in the ordering of these events throughout the simulations when comparing the order of acquisition to the order of first-production (**Figure 2B**). Beyond the “seed” agent, we found large variation in these orders, with only 24.75% of agents producing the behaviour in the same order as acquiring knowledge of it (i.e. light diagonal pattern in **Figure 2B**), and a divergence score of *d*_order_ = 1.95[1.13, 2.71]. This can be interpreted as after agent *i* acquired knowledge of the novel behaviour, approximately 2 more agents acquired knowledge of it before agent *i* first produced it. Simulations obtained a mean delay score of *d*_time_ = 5.33[3.46, 7.13], representing the amount of time that passed between an agent acquiring a behaviour and producing it. Substantial divergence between orders of acquisition and production meant that the observed order of first-productions was not well correlated with the underlying network structure, as in **Figure 2**. This could cause errors in inference, which we return to in **Section 3.2**.

### 3.1 Production rules influenced tempo, divergence and delay

Our model yielded valuable insights into how rules that govern production choices influence diffusion dynamics. A sensitivity analysis showed that changes to almost any production parameter influenced diffusion dynamics, as measured by tempo, divergence and delay (further exploration of transmission parameters in **Section S2, Table S1**). Below we describe how varying the way agent’s made decisions influenced diffusion dynamics, detailed by parameter in **Table 2**.

**Table 2:**
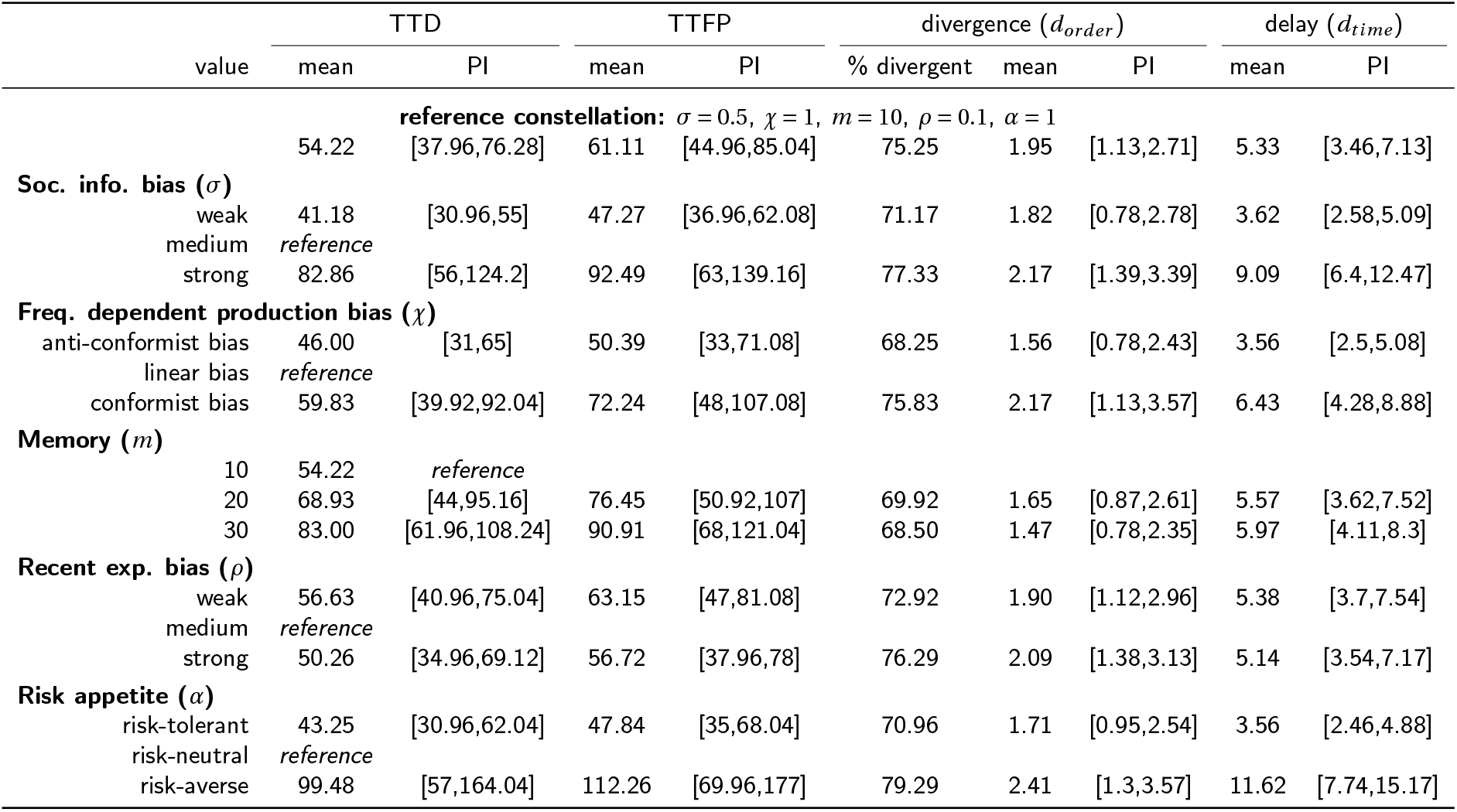
Summary of results. Mean and 92% percentile interval (PI) for time-to-diffusion, time-to-first-production, divergence of orders of acquisition and first-production, and time delay between acquisition and first-production. The reference constellation of parameter values is given first, and rows are arranged by model parameters in the order presented in the main text.

We first explored the effect of parameters that influenced how social information was used in production decisions. A weak social information bias (*σ*) resulted in a faster diffusion tempo with less divergence and delay compared to reference (**Equation** (8)). Conversely, a strong *σ* slowed diffusion (**Figure 3**, see **Figure S3** for all parameters), and increased divergence and delay. This effect was explained by the large increases in delay between acquisition and production (*d*_time_ in **Table 2**). When *σ* was strong, longer delays were obtained early in the diffusion, and when *σ* was weak, the delay remained consistent throughout the diffusion (**Figure S4A**). Agents with a weak *σ* were less influenced by the behaviour of associates, elevating the probability producing the novel behaviour upon acquisition and consequently hastening its diffusion (**Figure S5A**).

**Figure 3:**
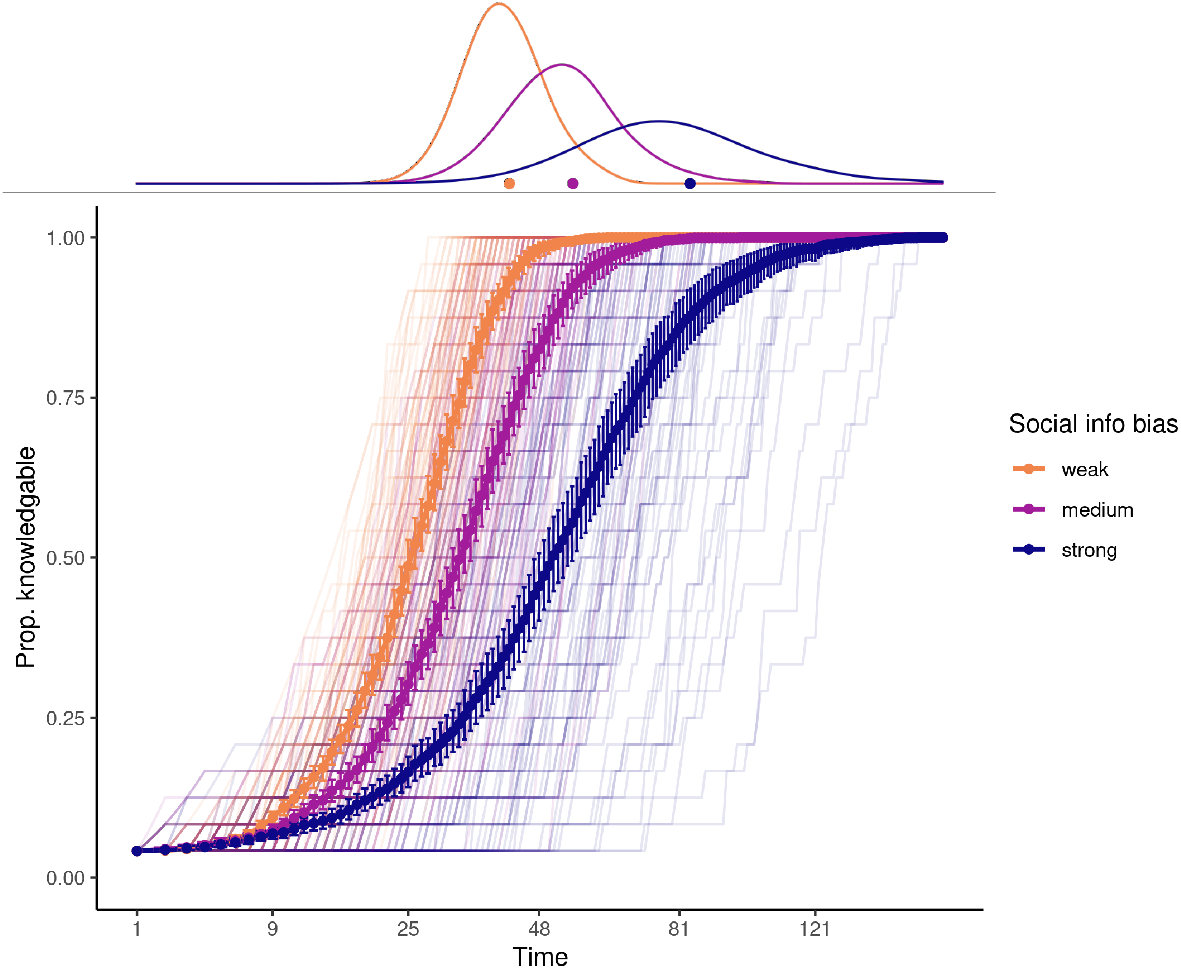
Strong social information bias slowed diffusion. Diffusion curves for the novel behaviour with distributions of time-to-diffusion in top margin, pooled within social information bias (colour, mean with 95% bootstrapped CI and traces from individual simulations). Data shown for reference level (300 simulations) and x-axis is square-root transformed for ease of reading. Strong social information bias within the sub-model of production greatly slowed diffusion.

We then explored the frequency dependent production bias. Compared to reference, production conformity (*χ* = 3) increased mean tempo, divergence and delay, while anti-conformity resulted in the opposite effects (**Table 2, Figure S3**). Conformity consistently increased delay relative to intervals between acquisition events throughout diffusion (**Figure S4B**). Conformity amplified the social influence of naive associates, causing agents to prefer the established behaviour longer after acquisition (**Figure S5B**). This delayed the learning of the expected payoff of the novel behaviour, lowered production rates, and thus lowered the probability of further transmission.

Finally, relevant for both production and acquisition sub-models was the memory window (*m*). Increasing agent’s memory slowed diffusion, yet resulted in less overall divergence (**Table 2, Figure S3B**). This was because larger memory slightly increased delays, yet generated more consistent delay intervals throughout diffusion, while increasing intervals between acquisition events (**Figure S4C**). With a large memory, there was greater cultural inertia, as any single observation of the novel behaviour meant less to an agent, as it was outweighed by memories of many observations of the established behaviour (**Figure S5C**).

Next, we considered parameters relevant for personal experience. A strong bias towards recent experience (*ρ*) resulted in a faster tempo than reference, and reduced divergence and delay (**Table 2, Figure S3C**). Larger *ρ* values hastened learning of expected payoffs of the novel behaviour, once acquired (**Figure S5D**). This made the novel behaviour more competitive against the established behaviour, as agents would be as likely to produce one as the other once their expected payoffs were similar.

Finally, risk-averse agents (*α* = .5) greatly slowed tempo, increased divergence, and nearly doubled delay (**Table 2, Figure S3D**) while risk-tolerant agents (*α* = 2) showed the reverse effects. In populations of risk-averse agents, delay decreased as the diffusion progressed as the productions of the novel behaviour by a growing number knowledgeable agents offset the effect of risk-aversion (**Figure S4E**). Risk-averse agents preferred the established behaviour for longer periods after acquisition, as past experience had built up their expectations of the established behaviour, while they had no expectations of the novel behaviour (**Figure S5E**). This decreased the competitiveness of the novel behaviour against the established behaviour.

In summary, changes in production rules that slowed diffusion (e.g. strong social information bias, conformity, risk-aversion) generally did so by decreasing the likelihood of production of the novel behaviour by newly knowledgeable agents. Any decrease in the likelihood of production was magnified by a feedback loop of reinforcement learning: lack of personal experience led to lower expectations of the novel behaviour, which encouraged the continued production of the established behaviour. Production rules that hastened diffusion (e.g. anti-conformity, strong bias towards recent experience and risk-tolerance) did so by increasing the relative competitiveness of the novel behaviour, once acquired.

### 3.2 The performance of inferential models when acquisition and production were linked

Continuous time-of-acquisition diffusion NBDA analyses (TADAc [14]) on “ideal” time-of-acquisition data yielded support for social transmission over asocial innovation as the acquisition mechanism for all social transmission simulations (median support Δ*AICc*: 20.5, **Figure S6A**). Under asocial innovation only, we found that support was ambivalent between asocial and social models (median Δ*AICc*: −2.38). The inferential models estimated the rate of social transmission close to that which had generated the data (**Figure S7A**). There was a trade-off between estimates for social transmission rate (*ŝ*) and base transmission rate (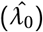), as simulation variance led to differences in how strictly the behaviour strictly followed the network. In summary, these results mirrored previous results regarding the inferential properties of NBDA applied to acquisition data.

We then tested TADAc on more realistic first-production data, generated through linking the sub-models described in **Sections 2.1 and 2.2**. Generally, TADAc had more difficulty inferring the true acquisition mechanism. Out of 2430 simulations where social transmission was the true acquisition mechanism, the models estimated 102 false-negatives (support for asocial innovation), and the distribution of support shifted towards asocial support (median Δ*AICc*: 8.65, **Figure S6A**). The strongest drivers of false negatives were production conformity, present in 86% of all false negatives, followed by a strong social information bias and risk aversion, both present in 61% (**Table S2**). Out of 2430 simulations of asocial innovation, the models estimated 1318 false-positives supporting social transmission instead, and the mean support shifted above 0 (median Δ*AICc*: 0.41). Strong social information bias and risk-aversion all increased the prevalence of false positives (**Table S2**). False support, positive or negative, was correlated with increased delays between acquisition and production, as well as more divergence between the order of acquisition and the order of first-production (**Figure S6B**).

EWA inference on “ideal” data, where all agents had the same repertoire over time, yielded point estimates of all parameters that were close to the true values used to generate the data. We then inferred parameter values from simulations where the novel behaviour either diffused via social transmission, and again where the novel behaviour was asocially innovated. Summary statistics of parameter posteriors from all three conditions are summarized in **Table S3**, and visualized per condition in **Figure S8**. In simulations of heterogeneous repertoires with social transmission or asocial innovation, social information bias (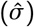) was over-estimated with the true value not contained in the 92% HPDI for 179 of 180 simulations (**Table S4**). Recent experience bias (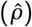) was overestimated and the true value was not contained in the 92% HPDI in most cases (105/180 estimates). These results show that heterogeneous knowledge states during diffusions negatively affect the ability of EWA to infer learning parameters. In particular, the overestimation of 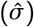 was an artifact of agents appearing to be influence by social information, simply because their knowledge state was constrained to the established behaviour.

## 4 Discussion

Understanding the relationship between psychological learning rules and population-level patterns is a long-standing research topic in cultural evolution [12, 35, 56, 57]. We contribute to this understanding by underlining the difference between learning rules that influence production *contra* acquisition. In doing so, we have uncovered new predictions for how changes to learning rules affect cultural diffusion dynamics by altering the relative competitiveness of novel behaviours. Our work highlights how learning during behavioural maintenance, a type of internal cultural selection, [9, 58, 59], should be considered another dimension that operates alongside external selection between individuals via transmission biases [60, 61, 62] and transformation of behaviour via cognitive biases [63, 64]. Not only does our model help to clarify discussions of strategies and biases, it has consequences for current practices in the field.

The case study of selective attrition in passerine birdsong illustrates why differentiating between acquisition and production matters [65, 66, 67, 68, 69]. Juvenile birds acquire song components from adult conspecifics, with an age-dependent transmission bias. However, as an adult, males preferentially produce the most frequently heard components, akin to a production conformity bias. This selective attribution then has consequences for repertoire size and composition (species with a fixed acquisition periods [65, 66, 67]; open-ended learners [68, 69]). Social learning strategies that occur either during acquisition or production have often been treated as interchangeable [10, 11, 12]; our model facilitates clear identification of where strategies or biases occur. For example, conformity could be a transmission bias, such as when naive great tits disproportionately acquire the majority produced solution to a foraging puzzle [20], or production bias, such as when a chimpanzees switch preference to match the group [70]. Our model allows for the explicit definition of combinations of biases acting on either process. These may yield similar effects, for example, both transmission [71] and production conformity slow diffusion. However, the effects of production biases are not necessarily predictable without linking acquisition and production, and more exploration is needed to determine whether a bias in one process might appear to be a different bias in another.

Our model assumed that in order to produce a behaviour, an individual must know that it can produce that behaviour, although it is not always easy to draw a line between acquisition and production. For example, many traits including birdsong, are acquired through repeated observation and practice. However, our sub-model of acquisition was ambivalent towards the precise mechanism of how individuals acquire this knowledge [8], and could represent indirect social learning (e.g. enhancement) or more direct social learning (e.g. observational conditioning) [5]. Our model could therefore be interpreted as accounting for acquisition via practice, assuming that practice productions did not influence associates or the expected payoffs of the practicing agent. Our model could be further extended by implementing more complicated acquisition mechanisms (e.g. complex contagion [72]) or a function to define how payoffs dynamically change with experience.

Our findings also have consequences for the way we analyse real-world data. We found that NBDA inference was less accurate in populations that were risk-averse, or highly sensitive to social information due to increased divergence between orders of acquisition and production. Therefore, NBDA is perhaps best suited to larger, sparser association networks, as longer intervals between acquisition events between clusters of nodes would help minimize divergence. We demonstrated that social information bias in EWA measures only social influence, and cannot distinguish between social or asocial acquisition of novel behaviour. Since the label of “social learning” has been given to both social transmission and social influence, this could lead those less familiar with EWA to misinterpret this parameter as evidence for social transmission. EWA might be of greater use when one can be relatively certain of homogeneous knowledge states. This includes the use of 1) experimental designs in which all choices are presented in a decision environment, 2) closed, rather than open-ended, tasks that constrain knowledge states and are less susceptible to morphological constraints, or 3) smaller social groups, where association is homogeneous and diffusion might occur rapidly relative to production rate.

In summary, we have shown how individual production decisions influence cultural diffusion dynamics at the population-level. We rendered a clear distinction between acquisition and expression of behaviour, and thus social learning and social influence. We argue this distinction improves definitions of social learning biases, and highlights the important role of reinforcement learning in explaining patterns of cultural change. Finally, our work encourages further development of methods to account for divergences between acquisition and production, which may be much more commonplace than previously acknowledged.

## 5 Acknowledgements

MC thanks Sonja Wild and Richard McElreath for their discussions. MC was supported by the IMPRS QBEE. MC, BJB, AK and LMA were supported by the Max Planck Society. MC, BJB, and LMA were supported by the CASCB, funded by the Deutsche Forschungsgemeinschaft (DFG) under Germany’s Excellence Strategy (EXC 2117-422037984).

## 6 Data Availability

Code and data for statistical analyses, figures, and agent based models are available at https://github.com/michaelchimento/acquisition_production_abm. A DOI will be assigned upon acceptance.

