## Supplementary materials for "Cultural diffusion dynamics depend on behavioural production rules"

#### Text

- **Text S1**: Sub-models in isolation
- **Text S2**: Sensitivity analysis of acquisition parameters

#### Tables

- **Table S1**: Summary of results for acquisition parameters.
- **Table S2**: Contributions of parameters to false negatives and positives in NBDA inference.
- **Table S3**: Posterior distributions from EWA inference on simulated data.
- **Table S4**: Inferential EWA performance by condition.

#### Figures

- **Figure S1**: Exemplar networks.
- **Figure S2**: Simulation dynamics of isolated production and acquisition sub-models.
- **Figure S3**: Diffusion curves for parameter values of production sub-model.
- **Figure S4**: Delay between acquisition and production against inter-acquisition intervals.
- **Figure S5**: Dynamics of internal states of agents following acquisition.
- **Figure S6**: Delay and divergence correlated with false support in NBDA analyses.
- **Figure S7**: Parameter estimates from NBDA analyses of simulated data.
- **Figure S8**: Posterior density estimates for parameters from EWA inference.

### S1 Sub-models in isolation

We explored either sub-model in isolation using the reference parameter constellation (**Equation 8**). Simulating data with the production sub-model alone required initialising all agents with knowledge of both behaviours, resulting in a flat acquisition curve constant at 100% of the population, and generating an r-shaped production curve that ended quickly (mean time-to-first-production [92% percentile interval]: $11.6\,[4.52, 20.5]$ time-steps; **Figure S2A**). We then simulated data where acquisition was not conditioned on production, which led to a knowledge acquisition curve that completed much faster than the first production curve (mean time-to-diffusion [PI] $5.70\,[4, 8]$ time-steps; **Figure S2B**). The shape of the production curve was not connected to the acquisition curve, leading to a growing distance between the two as the diffusion progressed (mean TTFP [PI]: $18.2\,[11, 27]$ time-steps). In summary, neither model alone was an adequate model of diffusion. In the isolated acquisition sub-model, production frequencies could not influence acquisition, and in the isolated production sub-model, agents must be initialized with both behaviors in repertoire.

### S2 Sensitivity analysis of acquisition parameters

Beyond manipulating the parameters related to production presented in the main text, we also varied four parameters related to the acquisition sub-model: network architecture, the baseline learning rate, the rate of social transmission ($s$) and the presence of asocial innovation, or innovation $A$.

In this section, we used the following parameter constellation as reference

$$\text{network} = \text{random regular}, \lambda_b = 0.05, s = 5, A = 0. \tag{1}$$

.

The parameters from the production sub-model were held at their reference value (main text **Equation 8**). The results of manipulating acquisition parameters are summarized in **Table S1**.

In addition to random regular networks, we tested Watts-Strogatz small world networks, Erdős–Rényi networks and Barabási–Albert scale free networks. Small world networks were fixed degree ($k = 6$), and ER and BA networks were parameterized such that $\bar{k} = 6$. ER networks are a type of low-clustering random network that created under the assumption that each link is independent and equally likely. It results in a graph that has a Poisson degree distribution. For each simulation, we generated networks until the resulting network contained 0 orphaned nodes. BA networks are random networks created under an assumption of preferential attachment of new nodes. Their degree distribution follows a scale-free power law. Degree was standardized between networks, as the probabilities generated by the sub-model of acquisition are degree dependent.

We found some variation in diffusion tempo and divergence between network types. While holding all other parameters at the reference level, random regular networks obtained the fastest tempo, followed by Erdős–Rényi networks, Barabási–Albert networks and small world networks (see **Table S1**, network architecture). We found that small-world networks obtained the slowest diffusion, and the smallest divergence score between orders of acquisition and production (59.8% lower mean divergence score compared to reference). However, the overall percentage of divergence agents was only reduced by 8%. These differences were likely due to the fact that we constructed small world networks without any probability of rewiring, leading to high local clustering within neighborhoods of agents ($C = 0.6$ when $K = 6$) and longer average path lengths. Neighborhoods of individual agents were thus uniform lattice structures arranged within a ring. The novel behaviour only had two directions from which to spread from the seed agent around the ring structure of the network, creating a longer, more uniform and predictable diffusion.

The social transmission rate $s$ influenced the probability of acquisition in a given time-step per unit connection to knowledgeable individuals, thus having consequences for diffusion tempo, as well as divergence in orders of production and transmission. Increasing the value of $s$ increased the tempo of diffusion as behaviours took fewer time-steps to spread from knowledgeable to naïve agents (see **Table S1**, $s$). More interestingly, it also increased amount of variation between order of acquisition and production, with 82% of agents obtaining divergent orders and a divergence score nearly 50% higher under the largest $s$ value tested, when compared with the reference setting. This was because the higher $s$ was, the more neighbours might have acquired knowledge of the novel behaviour before the next first-production event. Generally, longer periods between acquisition events lead to less divergence between orders of acquisition and production.

The baseline learning rate $\lambda_b$ influenced the probability of acquisition through either social transmission or asocial innovation by multiplying $s$ and $A$. At the reference level, asocial innovation was impossible ($A = 0$), so $\lambda_b$ only influenced social transmission, and thus was similar in effect to the direct manipulation of $s$, detailed below. Interestingly, even at very low probabilities of social transmission ($\lambda_b = 0.01$), 56.92% of agents still obtained divergent orders, although the divergence score was reduced (mean $d_{order} = 1.12\,[0.52, 2.05]$).

The presence of asocial innovation $A = 1$ halved the tempo of diffusion, as agents could acquire the behaviour independent of their associates' knowledge states. It also greatly increased the divergence (mean $d_{order} = 3.25\,[4.43, 11.57]$), while slightly decreasing delay (mean $d_{time} = 5.23\,[3.41, 7.30]$).

Table S1: **Summary of results for transmission parameters**. Mean and 92% PI values for time-to-diffusion (TTD), time-to-first-production (TTFP), divergence of orders of acquisition and first production ($d_{\text{order}}$), and time delay between acquisition and first production ($d_{\text{time}}$). The reference constellation of parameter values is given first, and rows are arranged by model parameters.

| | TTD | | TTFP | | divergence $d_{order}$ | | | delay $d_{time}$ | |
|---|---|---|---|---|---|---|---|---|---|
| value | mean | PI | mean | PI | % divergent | mean | PI | mean | PI |
| **reference constellation:** network=random regular, $s = 5$, $\lambda_b = 0.05$, $A = 0$ | | | | | | | | | |
| | 55.52 | [34,81.04] | 61.98 | [42.92,91.08] | 73.50 | 2.02 | [1.04,3.22] | 5.34 | [3.71,7.81] |
| **Network architecture** | | | | | | | | | |
| random regular | *reference* | | | | | | | | |
| small world | 75.19 | [51.88,104.32] | 80.95 | [57,116] | 65.46 | 1.22 | [0.69,1.83] | 5.19 | [3.38,7.04] |
| erdos | 58.63 | [34,90.12] | 64.54 | [37.92,96.04] | 71.71 | 1.85 | [1.04,2.7] | 5.11 | [3.16,7.08] |
| barabasi | 60.82 | [35,90.12] | 66.47 | [40,98.12] | 72.21 | 1.90 | [1.13,2.7] | 5.14 | [3.74,7.18] |
| **Social trans. rate ($s$)** | | | | | | | | | |
| 1 | 152.70 | [101,215.12] | 156.82 | [107.96,220.08] | 47.50 | 0.76 | [0.26,1.3] | 5.19 | [3.58,6.89] |
| 5 | | *reference* | | | | | | | |
| 10 | 38.78 | [25,60.2] | 47.05 | [30.96,70.32] | 80.38 | 2.67 | [1.74,3.83] | 5.25 | [3.5,7.54] |
| 15 | 31.34 | [21.96,46.08] | 40.38 | [30.96,58.12] | 82.25 | 3.11 | [2,4.63] | 5.46 | [3.82,6.96] |
| **Baseline learning rate ($\lambda_b$)** | | | | | | | | | |
| 0.01 | 150.66 | [95,215.44] | 155.37 | [98.96,222.24] | 56.92 | 1.12 | [0.52,2.05] | 5.28 | [3.33,7.17] |
| 0.05 | *reference* | | | | | | | | |
| 0.1 | 38.03 | [26.96,54.04] | 46.10 | [34,63.04] | 87.79 | 3.54 | [2.04,5.35] | 5.17 | [3.62,7] |
| **Asocial learning ($A$)** | | | | | | | | | |
| 0 | *reference* | | | | | | | | |
| 1 | 24.70 | [19,32.04] | 32.53 | [24.96,41] | 85.38 | 3.25 | [2,4.52] | 5.23 | [3.41,7.3] |

Table S2: **Contributions of parameters to false negatives and positives in NBDA inference**. Summary of the number of simulations (out of 2430) resulting in either false negative ($AICc_{asocial} - AICc_{social} \leq 0$ when data was generated with social transmission) or false positive support ($AICc_{asocial} - AICc_{social} > 0$ when data was generated with asocial innovation), according to relative AICc scores.

| value | false negatives | % sims w/ value | % all FN | false positives | % sims w/ value | % all FP |
|---|---|---|---|---|---|---|
| **Social info. bias** | | | | | | |
| weak | 16 | 1.98 | 15.69 | 340 | 41.98 | 25.80 |
| medium | 23 | 2.84 | 22.55 | 407 | 50.25 | 30.88 |
| strong | 63 | 7.78 | 61.76 | 571 | 70.49 | 43.32 |
| **Recent exp. bias** | | | | | | |
| weak | 30 | 3.70 | 29.41 | 424 | 52.35 | 32.17 |
| medium | 32 | 3.95 | 31.37 | 454 | 56.05 | 34.45 |
| strong | 40 | 4.94 | 39.22 | 440 | 54.32 | 33.38 |
| **Risk-appetite** | | | | | | |
| risk-tolerant | 22 | 2.72 | 21.57 | 329 | 40.62 | 24.96 |
| risk-neutral | 20 | 2.47 | 19.61 | 402 | 49.63 | 30.50 |
| risk-averse | 60 | 7.41 | 58.82 | 587 | 72.47 | 44.54 |
| **Freq. dependent production bias** | | | | | | |
| 0.33 | 1 | 0.12 | 0.98 | 446 | 55.06 | 33.84 |
| 1 | 13 | 1.60 | 12.75 | 486 | 60.00 | 36.87 |
| 3 | 88 | 10.86 | 86.27 | 386 | 47.65 | 29.29 |
| **Memory window** | | | | | | |
| 10 | 28 | 3.46 | 27.45 | 429 | 52.96 | 32.55 |
| 20 | 30 | 3.70 | 29.41 | 438 | 54.07 | 33.23 |
| 30 | 44 | 5.43 | 43.14 | 451 | 55.68 | 34.22 |

Table S3: **Posterior distributions from EWA inference on simulated data**. Posterior estimates for EWA parameters from three types of simulations (column headers), with each row containing the true parameter value used to generate data, the estimated mean and 92% HPDI, the mean and range of the Gelman-Rubin statistic ($\hat{R}$) and number of effective samples from the posterior across inference from all simulations containing the true value. In "homogeneous" simulations, all agents began with knowledge of both behaviours, without active diffusion of the novel behaviour. Here, parameter estimates were reasonably matched the true values used to simulate the data. In "heterogeneous social learning" simulations, agents began with knowledge of the establish behaviour, while a novel behaviour diffused via social learning. Recent experience bias ($\rho$) and social information bias ($\sigma$) were over-estimated. In "Heterogeneous asocial innovation" simulations, the behaviour diffused only through asocial innovation. Again, here $\rho$ and $\sigma$ are over-estimated.

| parameter | true value | estimate | rhat | effective sample size |
|---|---|---|---|---|
| **Homogenous** | | | | |
| **rho** | 0.25 | 0.3 [0.16,0.45] | 1.002 [0.999,1.006] | 1293 [1014,1562] |
| | 0.50 | 0.5 [0.3,0.67] | 1.002 [0.999,1.007] | 1255 [975,1500] |
| | 0.75 | 0.75 [0.61,0.88] | 1.002 [1,1.006] | 1229 [923,1511] |
| **sigma** | 0.25 | 0.27 [0.18,0.35] | 1.001 [1,1.006] | 1334 [935,1689] |
| | 0.50 | 0.5 [0.4,0.61] | 1.002 [0.999,1.008] | 1282 [966,1613] |
| | 0.75 | 0.75 [0.64,0.85] | 1.002 [0.999,1.005] | 1214 [897,1531] |
| **Heterogeneous (social transmission)** | | | | |
| **rho** | 0.25 | 0.6 [0.33,0.87] | 1.001 [0.999,1.005] | 1744 [1206,2095] |
| | 0.50 | 0.69 [0.45,0.93] | 1.001 [0.999,1.004] | 1771 [1300,2231] |
| | 0.75 | 0.8 [0.6,0.97] | 1 [0.999,1.003] | 1829 [1458,2375] |
| **sigma** | 0.25 | 0.65 [0.5,0.8] | 1.001 [0.999,1.003] | 1797 [1368,2482] |
| | 0.50 | 0.82 [0.71,0.92] | 1.001 [0.999,1.002] | 1907 [1482,2365] |
| | 0.75 | 0.93 [0.88,0.98] | 1 [0.999,1.006] | 1849 [1443,2204] |
| **Heterogeneous (asocial learning)** | | | | |
| **rho** | 0.25 | 0.52 [0.29,0.77] | 1.001 [0.999,1.003] | 1613 [1317,2004] |
| | 0.50 | 0.67 [0.48,0.89] | 1 [0.999,1.002] | 1670 [1308,2111] |
| | 0.75 | 0.81 [0.63,0.97] | 1.001 [0.999,1.004] | 1594 [1092,2074] |
| **sigma** | 0.25 | 0.52 [0.4,0.66] | 1.001 [0.999,1.002] | 1609 [1309,1935] |
| | 0.50 | 0.71 [0.6,0.81] | 1 [0.999,1.004] | 1713 [1331,2385] |
| | 0.75 | 0.86 [0.8,0.92] | 1.001 [0.999,1.004] | 1818 [1483,2285] |

Table S4: **Inferential EWA performance by condition**. For each combination of true values of $\rho$ and $\sigma$ (row), inference was run on 10 simulated data-sets. When agents had heterogeneous repertoires, the inferential ability of EWA was impaired, with especially $\hat{\sigma}$ consistently over-estimated. These results demonstrate that the use of EWA during diffusions should be avoided, as $\sigma$ is intended to represent social influence, and should not be taken to infer social transmission.

| True values | | true $\rho$ in HPDI | | true $\sigma$ in HPDI | |
|---|---|---|---|---|---|
| rho | sigma | # sims | % | # sims | % |
| **Homogeneous repertoires** | | | | | |
| 0.25 | 0.25 | 10 | 100 | 10 | 100 |
| 0.25 | 0.50 | 9 | 90 | 9 | 90 |
| 0.25 | 0.75 | 10 | 100 | 9 | 90 |
| 0.50 | 0.25 | 10 | 100 | 9 | 90 |
| 0.50 | 0.50 | 10 | 100 | 10 | 100 |
| 0.50 | 0.75 | 9 | 90 | 9 | 90 |
| 0.75 | 0.25 | 10 | 100 | 10 | 100 |
| 0.75 | 0.50 | 9 | 90 | 10 | 100 |
| 0.75 | 0.75 | 10 | 100 | 10 | 100 |
| **Heterogeneous repertoires (social transmission)** | | | | | |
| 0.25 | 0.25 | 0 | 0 | 0 | 0 |
| 0.25 | 0.50 | 0 | 0 | 0 | 0 |
| 0.25 | 0.75 | 2 | 20 | 0 | 0 |
| 0.50 | 0.25 | 0 | 0 | 0 | 0 |
| 0.50 | 0.50 | 4 | 40 | 0 | 0 |
| 0.50 | 0.75 | 9 | 90 | 0 | 0 |
| 0.75 | 0.25 | 5 | 50 | 0 | 0 |
| 0.75 | 0.50 | 10 | 100 | 0 | 0 |
| 0.75 | 0.75 | 10 | 100 | 0 | 0 |
| **Heterogeneous repertoires (asocial innovation)** | | | | | |
| 0.25 | 0.25 | 0 | 0 | 0 | 0 |
| 0.25 | 0.50 | 2 | 20 | 0 | 0 |
| 0.25 | 0.75 | 2 | 20 | 0 | 0 |
| 0.50 | 0.25 | 2 | 20 | 0 | 0 |
| 0.50 | 0.50 | 4 | 40 | 0 | 0 |
| 0.50 | 0.75 | 8 | 80 | 1 | 10 |
| 0.75 | 0.25 | 1 | 10 | 0 | 0 |
| 0.75 | 0.50 | 6 | 60 | 0 | 0 |
| 0.75 | 0.75 | 10 | 100 | 0 | 0 |

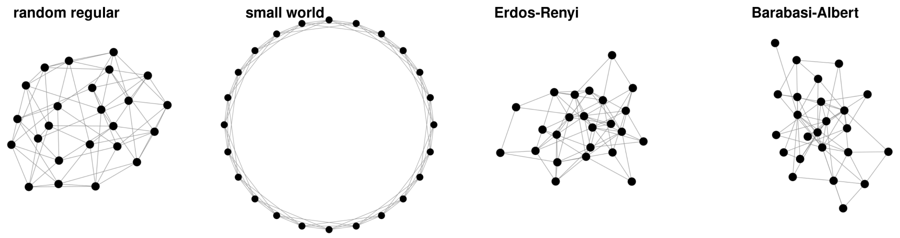


Figure S1: **Exemplar networks.** Four types of network architectures used for simulations. Random regular and small world networks contain nodes which each have 6 connections to neighbours. Erdős–Rényi and Barabási–Albert networks were parameterized such that the mean degree was 6 to make results comparable.

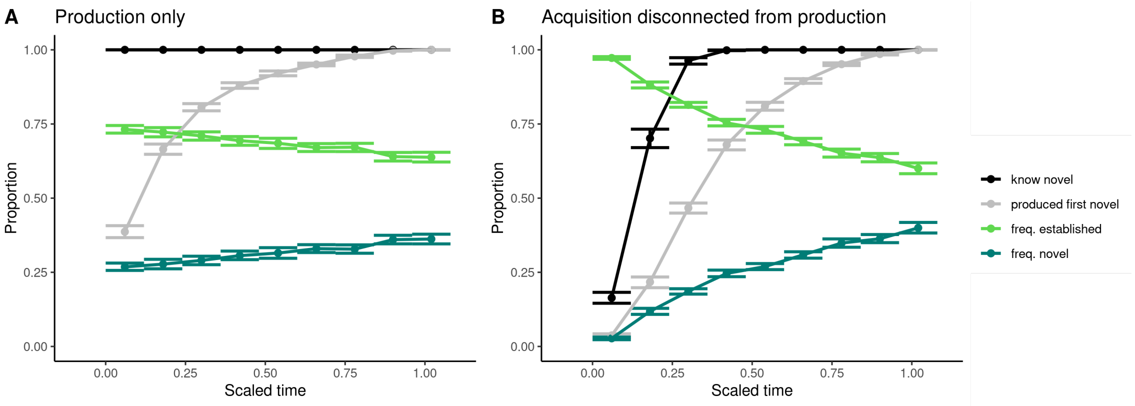


Figure S2: **Simulation dynamics of isolated production and acquisition sub-models.** Mean values (with 95% bootstrapped CI) over time (scaled between 0 and 1) of the proportion of agents which know the novel behaviour (black), the proportion of agents which have produced the novel behaviour at least once (grey), the proportion of behaviours that are the established behaviour (light green), and the novel behaviour (dark green). Parameters set at reference level (N=100 sims for each panel). **A)** Generating data with the sub-model of production only results in an r-shaped curve of first production, with approximately a quarter of all agents choosing to produce the novel behavior in the first time-step, despite having no prior experience with it, and no expected payoffs. Thus, the production sub-model is not appropriate for approximating diffusion as it greatly overestimates initial probabilities of production when most agents should not know the novel behavior, and produces no true acquisition curve. **B)** Generating data with the sub-model of transmission disconnected from production results in a diffusion curve that is completely disconnected from the first production curve, as agents can learn the novel behaviour without ever having seen it produced. The first production curve is only constrained by the transmission curve very early in the simulation, and more resembles the r-shaped curve in panel A.

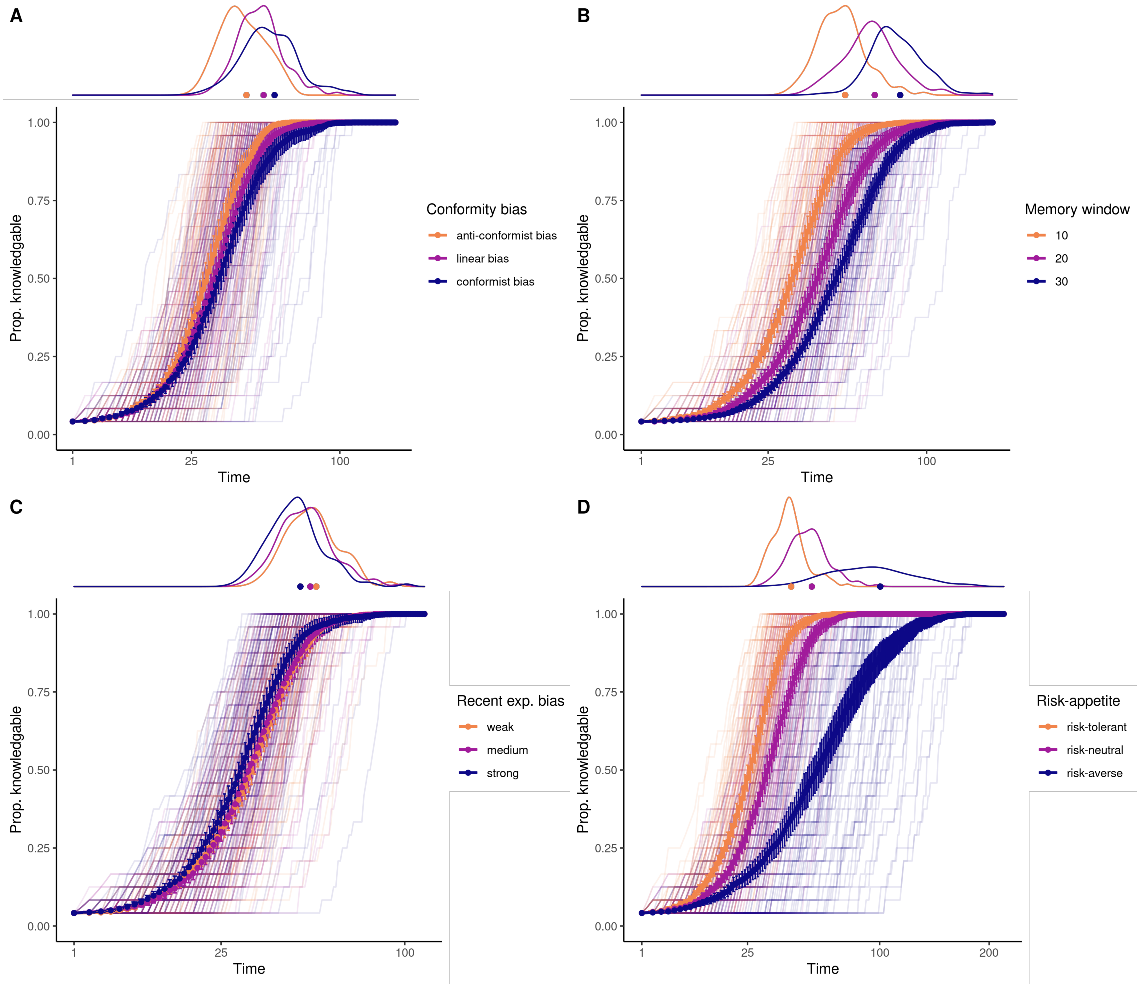


Figure S3: **Diffusion curves for parameter values of production sub-model.** Diffusion curves for the novel behaviour with distributions of time-to-diffusion in top margin, pooled within parameter value (colour, mean with 95% bootstrapped CI and traces from individual simulations). For each parameter, data shown holding all other parameters at reference level (300 simulations) and x-axis is square-root transformed for ease of reading.

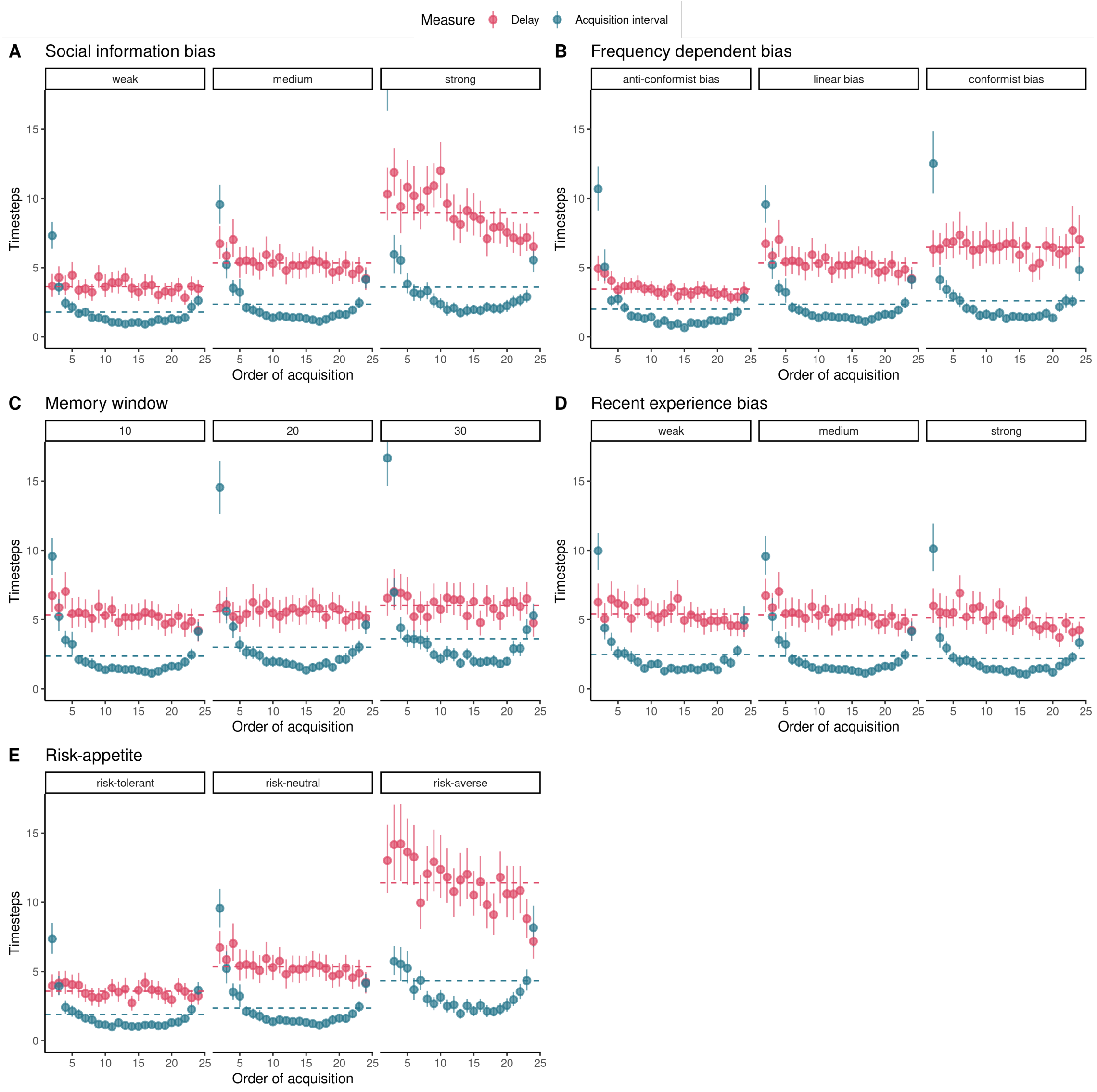


Figure S4: **Delay between acquisition and production against inter-acquisition intervals.** For each parameter manipulated in the main text, we plot the mean delay between acquisition and production events within an individual (red color), and the mean interval of time between acquisition events. We observed a decreasing relationship between order of acquisition and delays when social information bias was strong, and when agents were risk averse. Delays remained relatively constant over the order of acquisition when agents were conformist, or had larger memories.

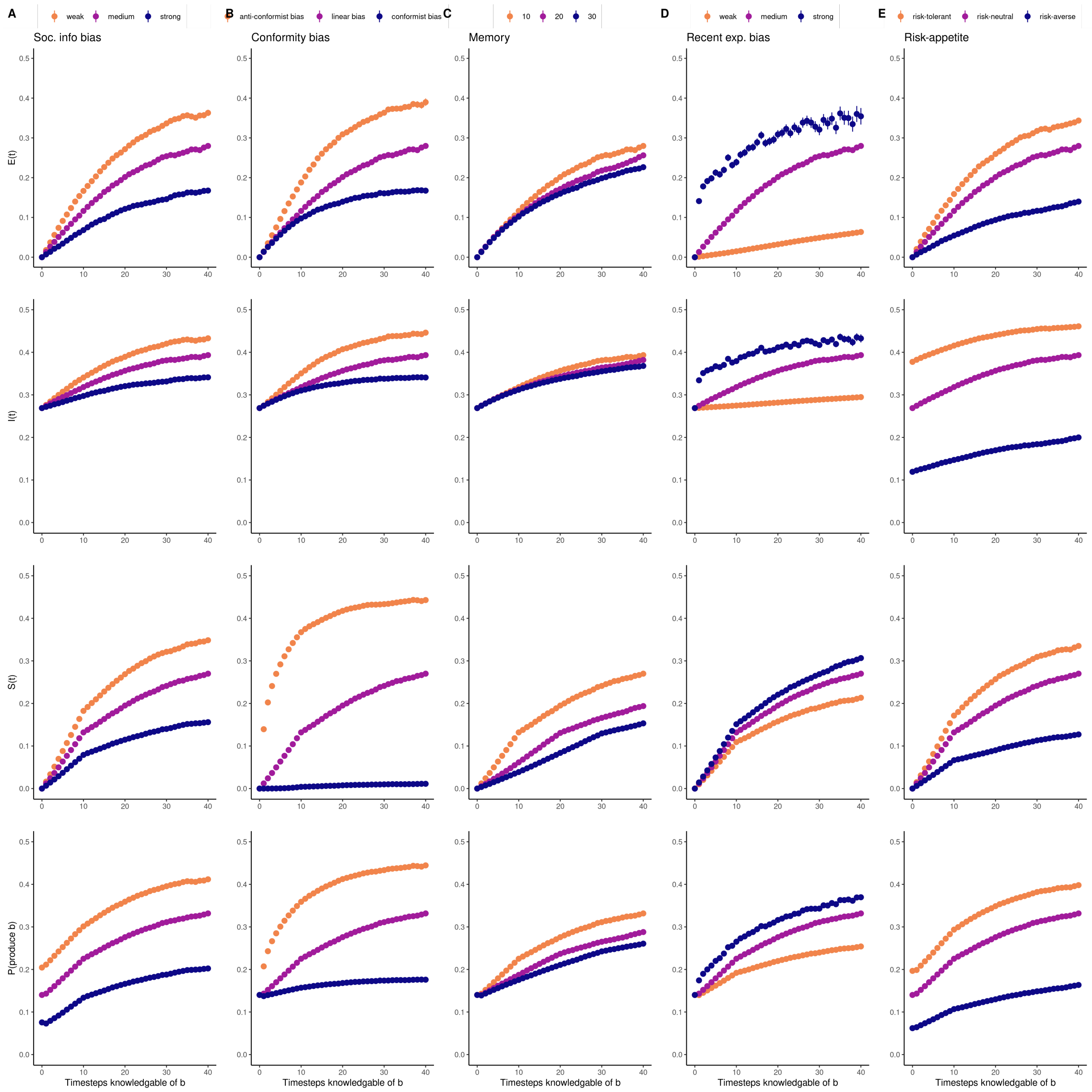


Figure S5: **Dynamics of internal states of agents following acquisition.** For each parameter manipulated in the main text, we plot the mean values of the expected payoff of novel behaviour $b$, its softmax transform, social observation values, and the final probability of production choice over the first 40 time-steps after agents acquired $b$.

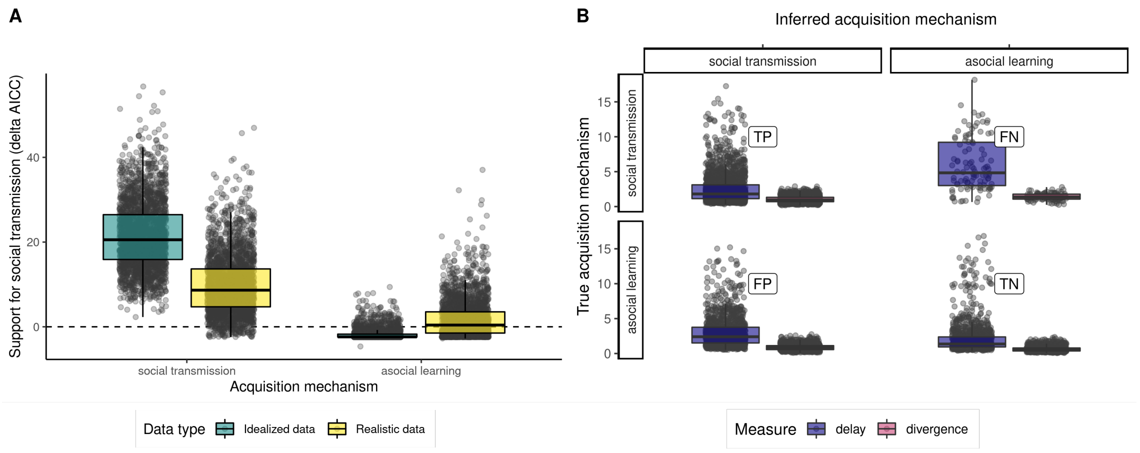


Figure S6: **Delay and divergence correlated with false support in NBDA analyses. A)** Comparison of relative support for social transmission ($\Delta AICc$) for idealized data (teal) and realistic data (yellow). Inference on idealized data matched our expectations, resulting in strong support for social transmission when social transmission was the generative mechanism behind the data, and support for asocial innovation when asocial innovation was the generative mechanism. However, using more realistic data, where transmission and production were linked, false support for the incorrect generative mechanism was more prevalent. **B)** We compared delay and divergence measures from each simulation among true and false positives and negatives (label). Simulations that received the correct support had lower delay and divergence scores, compared to false positives and negatives.

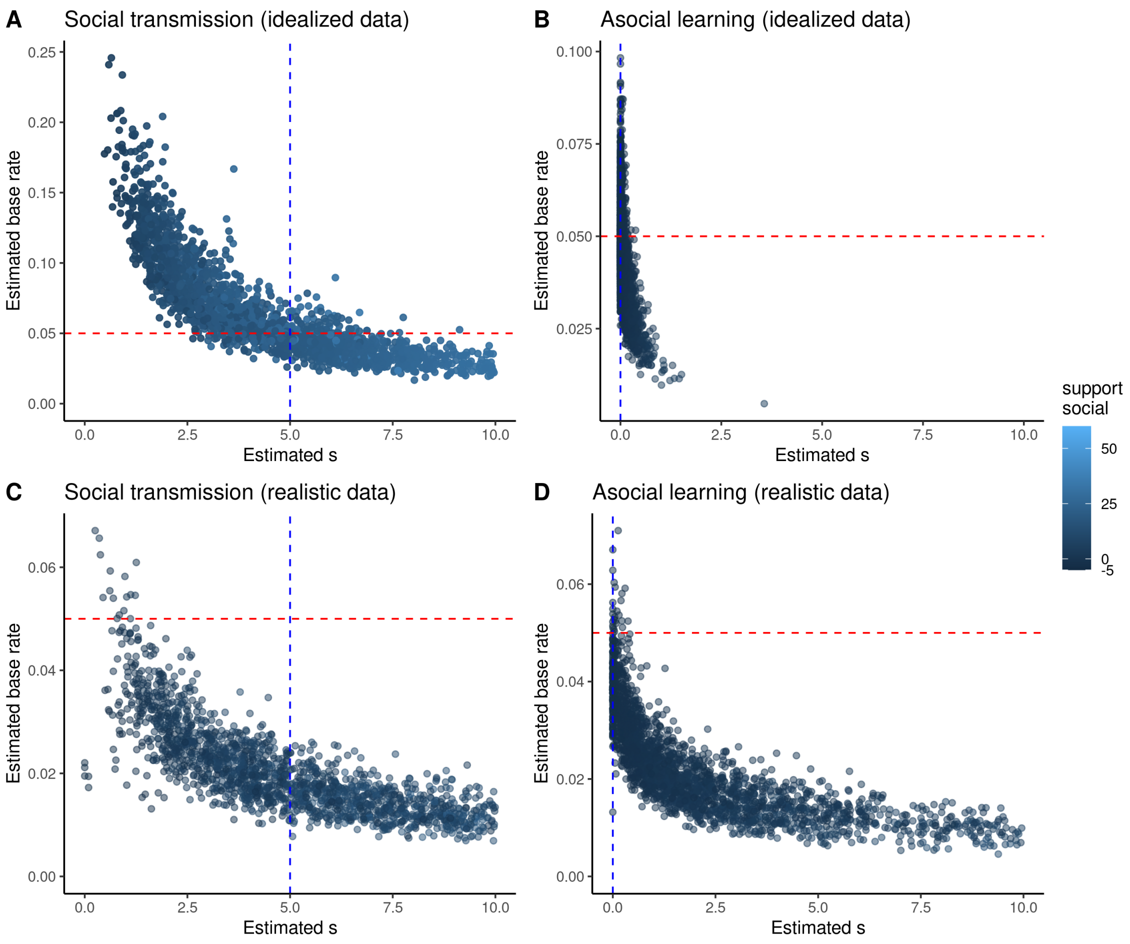


Figure S7: **Parameter estimates from NBDA analyses of simulated data** Each point represents parameter estimates for one simulation: rate of social transmission (x-axis) and baseline transmission rate values (y-axis). colour represents the support given to social transmission ($\Delta AICc$). Dashed lines represent true values of parameters used to generate data. **A)** Ideal data from a social transmission results in correct inference, although simulation variance leads to the curved shape of parameter estimates. If the diffusion follows the network closely, estimated strength of social transmission is inflated. **B)** Idealized asocial innovation data also results in correct inference, with very low s estimates and little support for social transmission. **C & D)** Realistic data is generated with transmission and production sub-models interacting. Parameters are mis-estimated in both cases, with many more false negative and positives than the idealized cases.

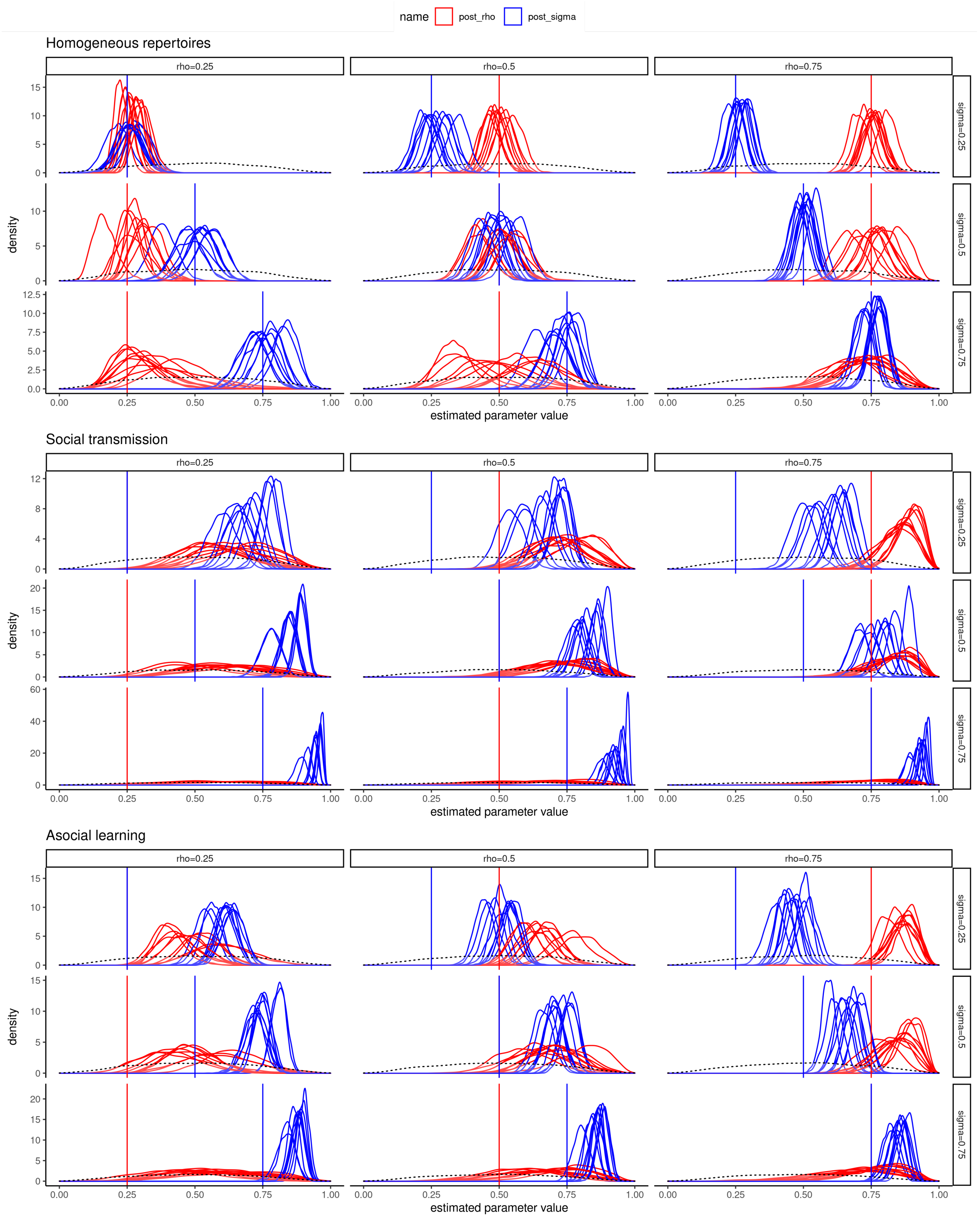


Figure S8: **Posterior density estimates for parameters from EWA inference.** Each panel shows the inferred posterior density estimates for recent experience bias ($\rho$) and social information bias ($\sigma$) for one simulation (N=1500 draws from posterior per parameter). The distribution with the dashed line represents the prior for both variables. True values used to simulate the data are shown with vertical lines, coloured by parameter. Inference on data simulated with homogeneous repertoires results in parameter estimates consistent with true values. Inference on data simulated with social transmission or asocial innovation results in general over estimation of both parameters.
